## Supplemental Figure for "Task-specific invariant representation in auditory cortex"

*Charles R. Heller*[1], *Gregory R. Hamersky*[1], *Stephen V. David*[2*]
[1]Neuroscience Graduate Program, [2]Otolaryngology, Oregon Health & Science University, Portland, OR, USA

***Supplemental materials***

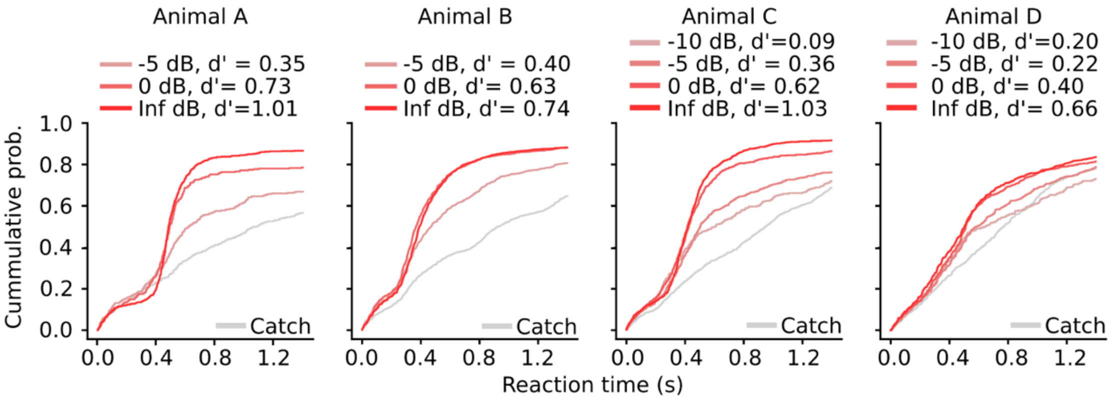


**Figure 1 – figure supplement 1. Behavioral performance of individual animals.** Cumulative reaction time histogram for each animal and target sound across all behavior sessions. Color indicates sound identity. Animals were rewarded for responses to all sounds except the catch.

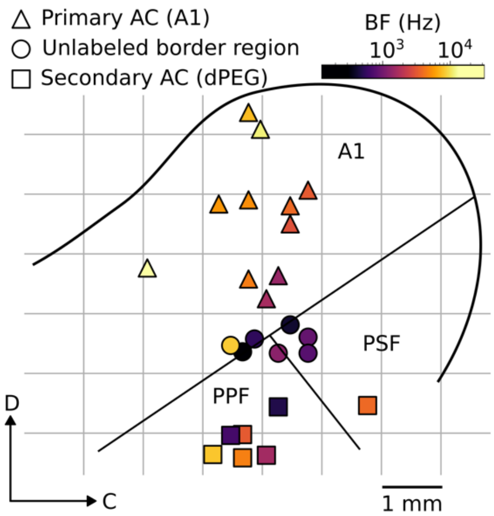


**Figure 2 – figure supplement 1. Penetration map for one example animal.** Left hemisphere from one animal. Each marker indicates the anatomical location of a single electrode penetration. Color indicates the average best frequency (BF) of neurons recorded at each location and marker type indicates which brain region each penetration belongs to. Overlaid black lines represent rough estimate of region boundaries. A1: Primary auditory cortex, PSF/PPF: Anterior / posterior fields of secondary auditory cortex (dPEG).

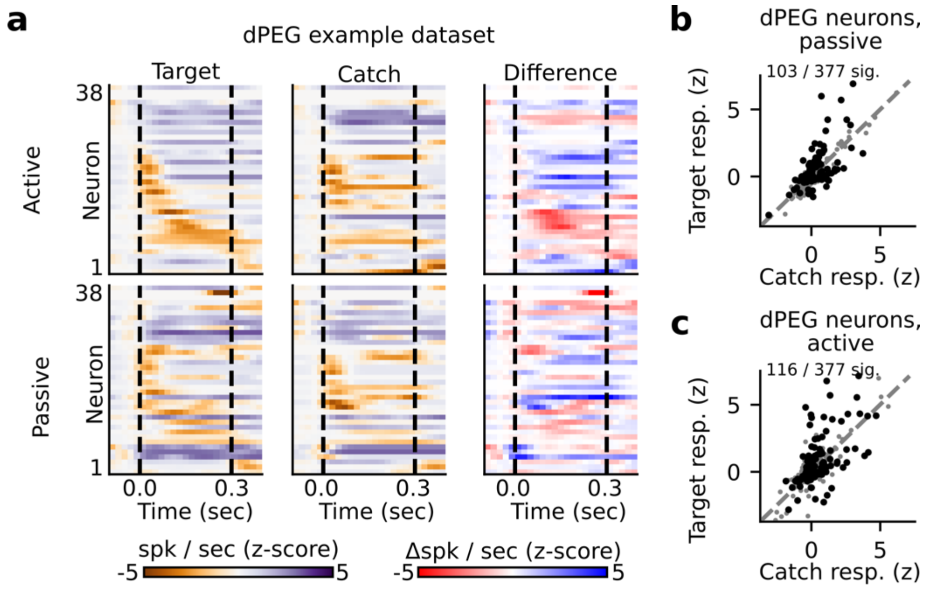


**Figure 2 – figure supplement 2. State-dependent modulation of singe neuron target vs. catch discrimination in dPEG. a.** Example peristimulus time histogram (PSTH) responses from a single recording site in dPEG. Heatmap color in each row indicates PSTH amplitude of one neuron. Dashed lines indicate sound onset / offset. Spikes were binned (20 ms), z-scored, and smoothed (30-ms Gaussian kernel). Example target responses are to the pure tone (Inf dB) target. Difference is computed as the z-scored response to the target minus the z-scored catch response (resulting in a difference shown in units of z-score). **b-c.** Mean z-scored response evoked by catch vs. Inf dB stimulus for each dPEG neuron across passive (**b**) and active (**c**) trials. Responses were defined as the total number of spikes recorded during the 300 ms of sound presentation (area between dashed lines in panel A). Neurons with a significantly different response to the catch vs. target stimulus are indicated in black and quantified on the respective figure panel.

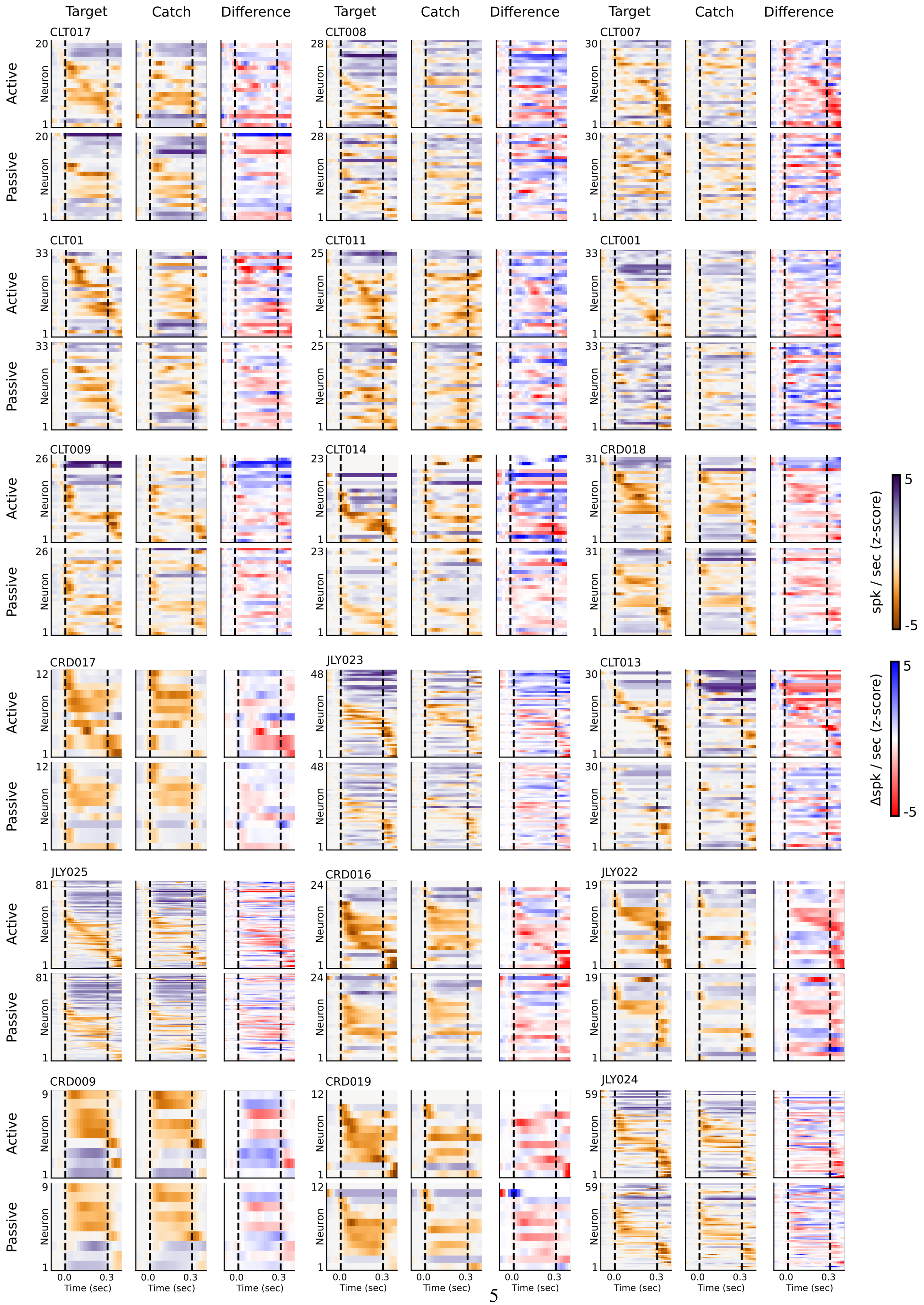

Target
Catch
Difference
Active
Passive
Neuron
CLT017
CLT008
CLT007
CLT01
CLT011
CLT001
CLT009
CLT014
CRD018
CRD017
JLY023
CLT013
JLY025
CRD016
JLY022
CRD009
CRD019
JLY024
Time (sec)
spk / sec (z-score)
Δspk / sec (z-score)

**Figure 2 – figure supplement 3. Single neuron target vs. catch raster plots for all A1 recording sites.** Example peristimulus time histogram (PSTH) responses from all recording sites in A1. Each group of six panels corresponds to a single recording site, as in Figure 2 and Supplemental Figure 3. Heatmap color in each row of each panel indicates PSTH amplitude of one neuron. Dashed lines indicate sound onset / offset. Spikes were binned (20 ms), z-scored, and smoothed (30-ms Gaussian kernel). Example target responses are to the pure tone (Inf dB) target. Difference is computed as the z-scored response to the target minus the z-scored catch response (resulting in a difference shown in units of z-score).

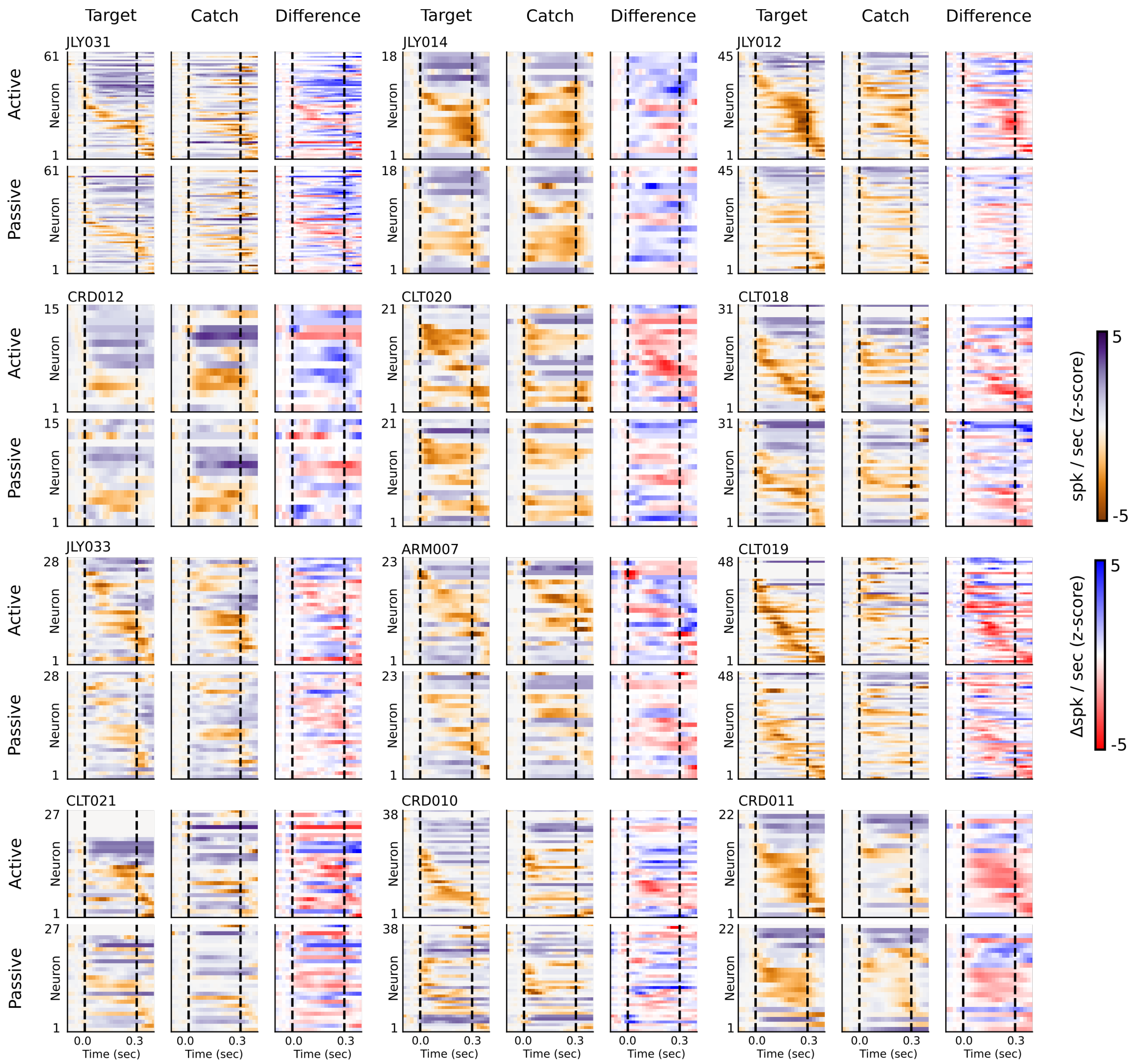


**Figure 2 – figure supplement 4. Single neuron target vs. catch raster plots for all dPEGrecording sites.** Example peristimulus time histogram (PSTH) responses from all recording sites in dPEG. Each group of six panels corresponds to a single recording site, as in Figure 2 and Supplemental Figure 3. Heatmap color in each row of each panel indicates PSTH amplitude of one neuron. Dashed lines indicate sound onset / offset. Spikes were binned (20 ms), z-scored, and smoothed (30-ms Gaussian kernel). Example target responses are to the pure tone (Inf dB) target. Difference is computed as the z-scored response to the target minus the z-scored catch response (resulting in a difference shown in units of z-score).

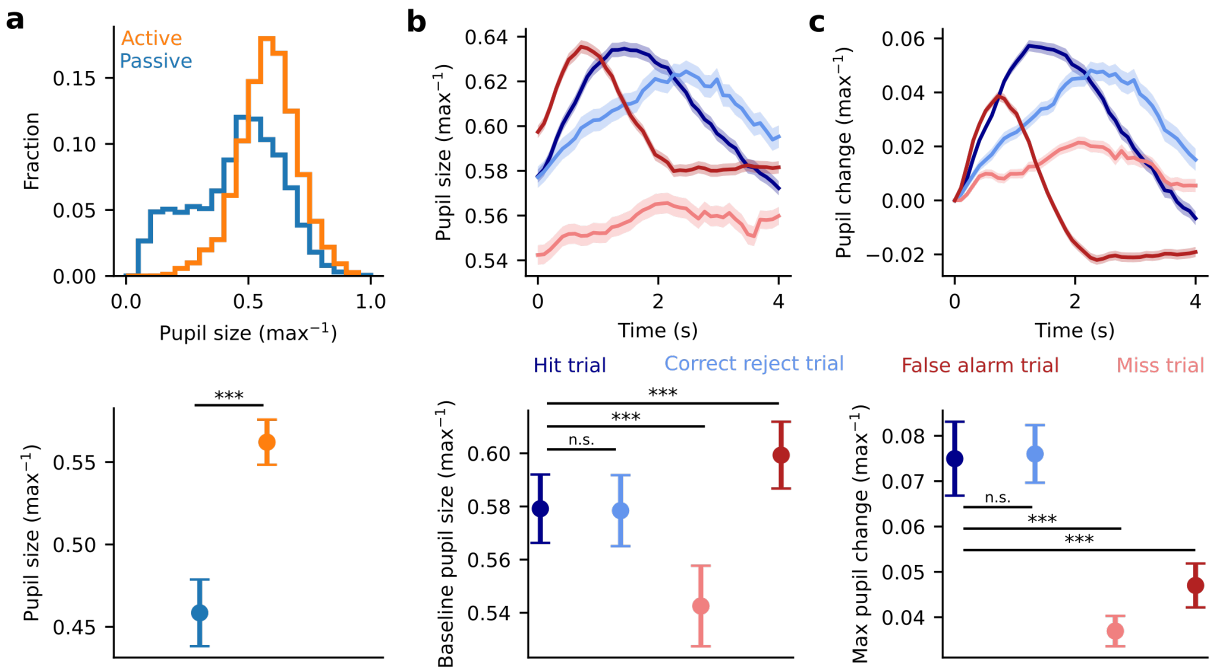


**Figure 3 – figure supplement 1. Pupil dynamics reflect both generalized arousal as well as trial outcome. a.** Top: Histogram of pupil size across all experiments (n = 39 sessions, n = 4 animals) during active (orange) and passive (blue) trials. Pupil size is normalized to the max pupil size observed within each experiment. Bottom: Mean pupil size +/- the standard error under each condition. Data were first averaged within each experiment before computing the mean, standard error across experiments, and before performing significance test. Pupil size was significantly larger on active trials than on passive trials (p = 1.48e-6, Wilcoxon signed rank test) **b**. Top: Raw, trial-weighted average of per-trial pupil on hit (dark blue), correct reject (light blue), false alarm (dark red), and miss (light red) trials. Bottom: Summary of mean pre-trial pupil for each type of trial outcome. Data were first averaged within each experiment before computing the mean, the standard error, and before performing significance tests (hit vs. correct reject p = 0.36, hit vs. miss p = 8.81e-5, hit vs. false alarm p = 2.83e-5, Wilcoxon signed rank test). **c**. Top: Same as in **b** after first normalizing pupil size to the pretrial mean. Bottom: Summary of the max change in pupil size during each trial for each type of trial outcome. Data were first averaged within each experiment before computing the mean, the standard error, and before performing significance tests (hit vs. correct reject p = 0.29, hit vs. miss p = 3.01e-5, hit vs. false alarm p = 1.62e-5, Wilcoxon signed rank test).

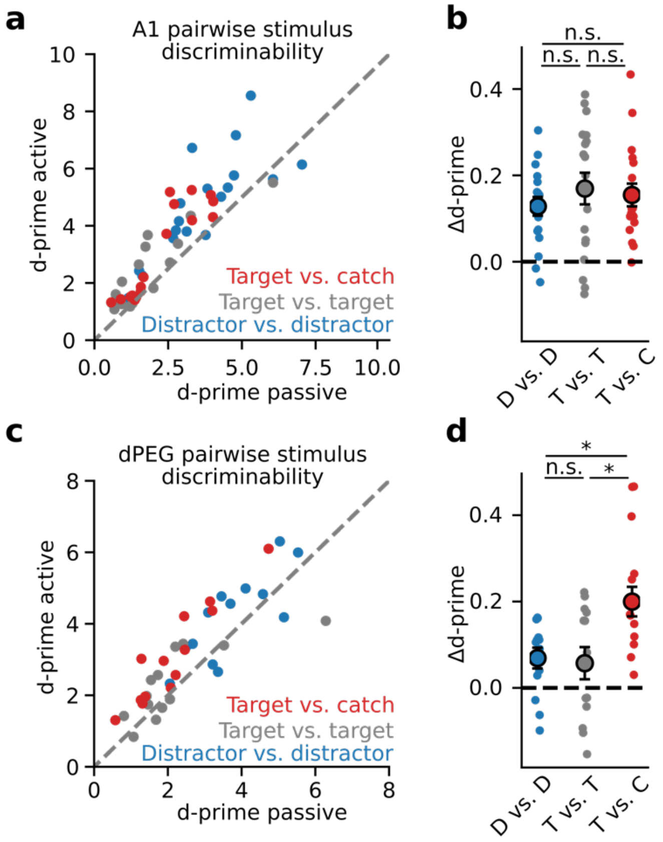


**Figure 3 – figure supplement 2. Selective enhancement of task-relevant category representation in secondary auditory cortex is not affected by global arousal.** Same as in Figure 3, without first correcting for pupil-indexed arousal explainable variance (*i.e.,* decoding stimulus identity from raw neural activity, without controlling for global arousal). **a.** Mean population d-prime between sounds from each category (target vs. catch, target vs. target, and distractor vs. distractor, Figure 1C) for each A1 recording site (n = 18 sessions, n = 3 animals). **b.** Δd-prime is the difference between active and passive d-prime, normalized by their sum (no significant difference, Wilcoxon signed rank test). **c.** Passive vs. Active category discriminability for dPEG recording sites, plotted as in A (n = 13 sessions, n = 4 animals). **d.** Changes in discriminability per category in dPEG. Δd-prime for target vs. catch pairs (T vs. C) was significantly greater than for the other categories (D vs. D: $p = 0.013$; T vs. T: $p = 0.017$, Wilcoxon signed rank test).

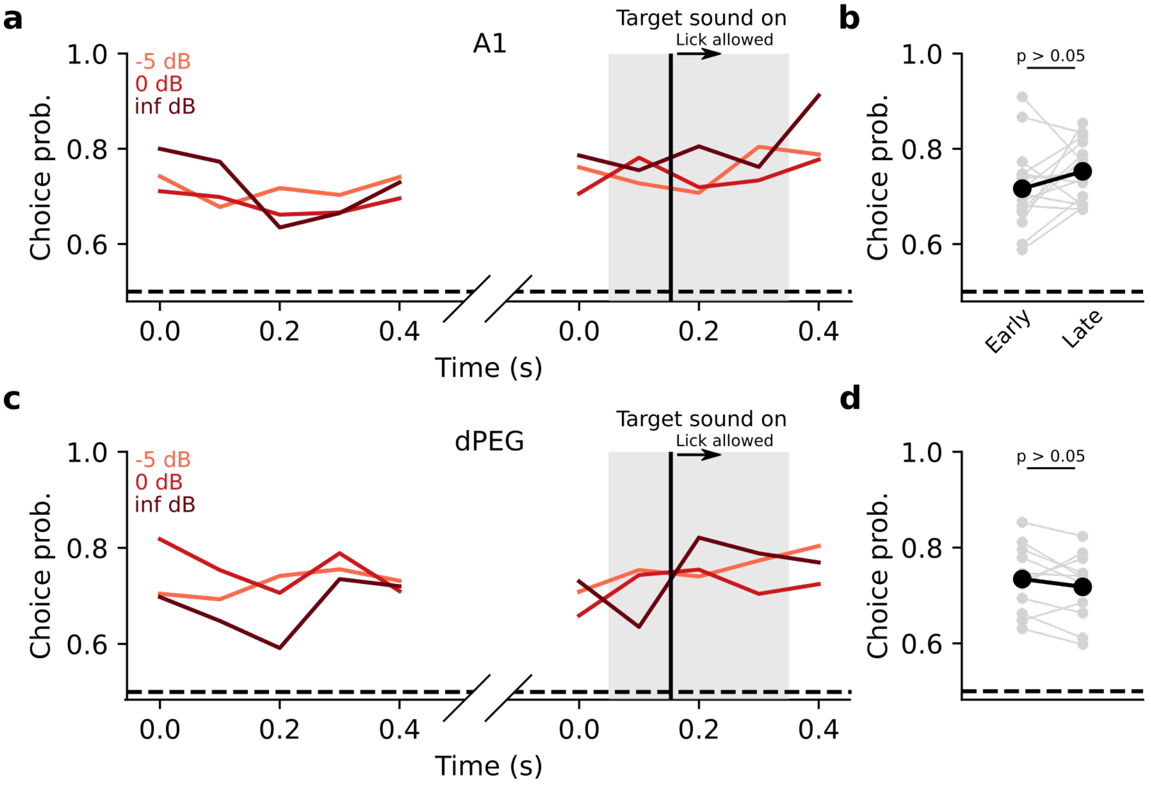


**Figure 4 – figure supplement 1. Choice decoding in auditory cortex primarily reflects impulsivity. a.** Mean choice probability on hit vs. miss trials across all A1 experiments, split by target stimulus SNR. Grey shading represents the time when the target sound stimulus is on. The vertical black line indicates when animals could begin licking in response to the target sound. Choice decoding is shown for the first distractor sound presentation on each trial (curves on the left) and for the target sound presentation (curves on the right). **b**. Comparison of mean choice probability during the first distractor sound (Early) to the mean choice probability during the target sound (Late). Each grey line represents one experiment, and the black line is the mean across experiments. Only bins prior to the lick window onset were analyzed to avoid motor activity confounds. There was no significant difference between early and late choice probability ($p > 0.05$, Wilcoxon signed rank test). **c-d.** Same as a-b for dPEG experiments.

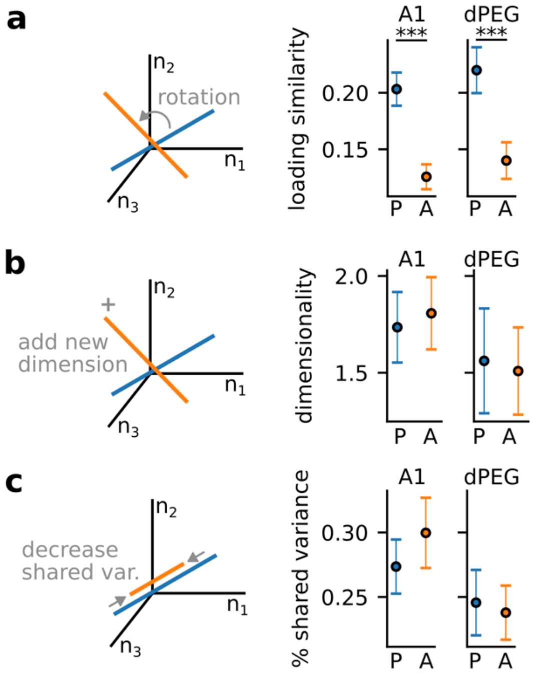


**Figure 5 – figure supplement 1. Modulation of population covariability metrics by task-engagement.** Population metrics extracted from Factor Analysis model fit. Models were fit to the catch stimulus only to control for variability due to stimulus identity. P=passive, A=active. **a.** Loading similarity is defined as the similarity of the neuronal loading weights onto the primary covariability axis (largest variance factor). Ranges from 0 (dissimilar) to 1 (similar). Loading similarity was significantly larger during passive conditions for both brain regions (A1: p = 6.8e-5, n = 18 sessions / 3 animals; dPEG: p = 6.9e-5, n = 13 sessions / 4 animals; Wilcoxon signed rank test). **b.** Dimensionality is defined as the number of significant dimensions of shared variability in the data determined by log-likelihood. No significant change in dimensionality was observed in either brain region. **c.** % shared variance is defined as the percentage of single neuron variance that can be explained by the shared, covariability axes. Ranges from 0 to 1. No significant change in % shared variance was observed in either brain region.

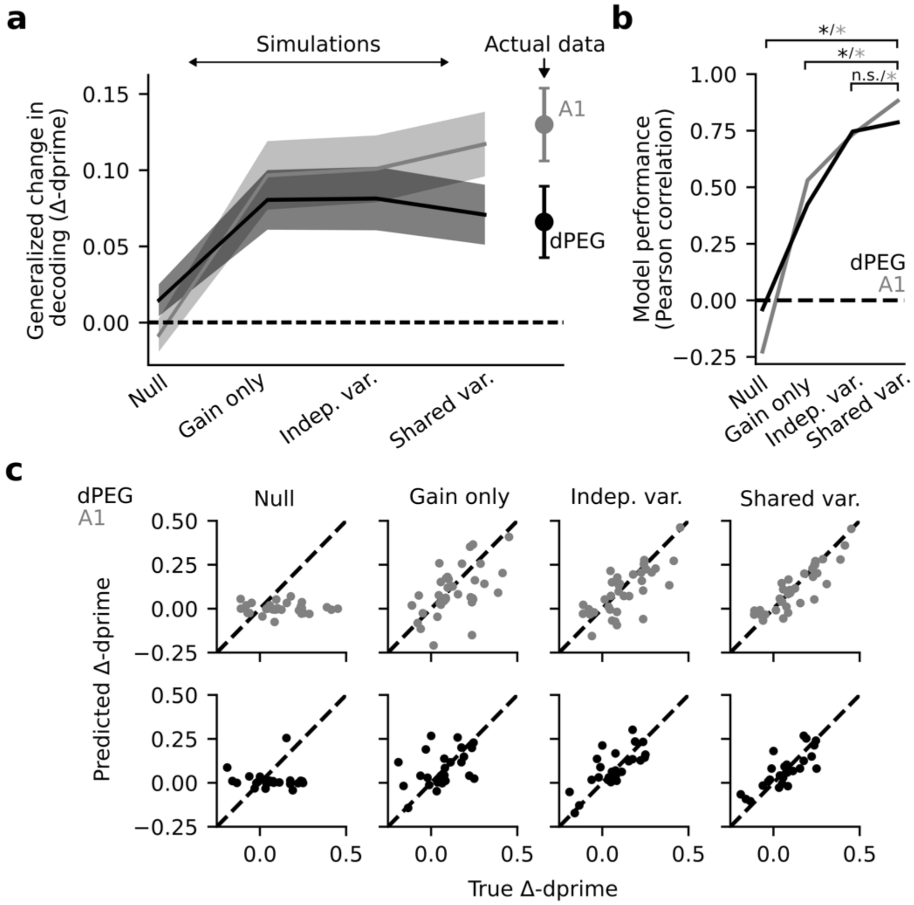


**Figure 5 – figure supplement 2. Modeling task-dependent changes in shared population covariability improve predictions of decoding changes in A1, but not dPEG. a.** Mean Δd-prime across all target vs. target and target vs. catch stimulus pairs for each Factor Analysis based simulation: Null model, gain only model, independent variance model, and shared variance model. Actual Δd-prime is shown on the right. **b.** Model performance is measured as the Pearson correlation between actual and predicted Δd-prime. In A1, model performance increases monotonically. In dPEG, including gain and independent variance both improve predictions but there is no significant improvement when including shared population covariability. Significance measured using bootstrap test with alpha = 0.05. **c.** Scatter plots show predicted vs. actual Δd-prime for each Factor Analysis simulation in both dPEG (black) and A1 (grey). Model performance (shown in B) was measured as the Pearson correlation of these scatter plots.
